## Supplemental figures for "BMP signaling maintains auricular chondrocyte identity and prevents microtia development by inhibiting protein kinase A"

**This file includes:**

**Supplemental figures**

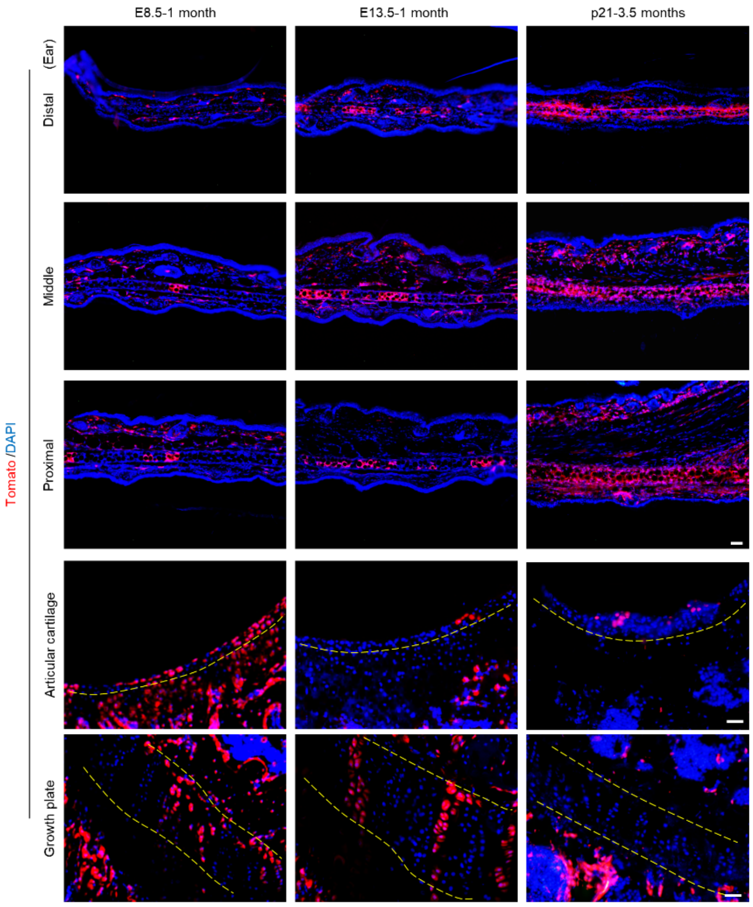


Figure 1-figure supplement 1

**Figure 1-figure supplement 1. Tracing of chondrocytes in auricle, articular cartilage, and growth plate in *Prrx1-CreERT; Bmpr1a*$^{f/f}$ mice.**
The mice received TAM at E8.5, E13.5, or p21 and the tissues were collected at 1 month or 3.5 months of age. Scale bars (ear)=100 µm. Scale bars (articular cartilage and growth plate) =50 µm.

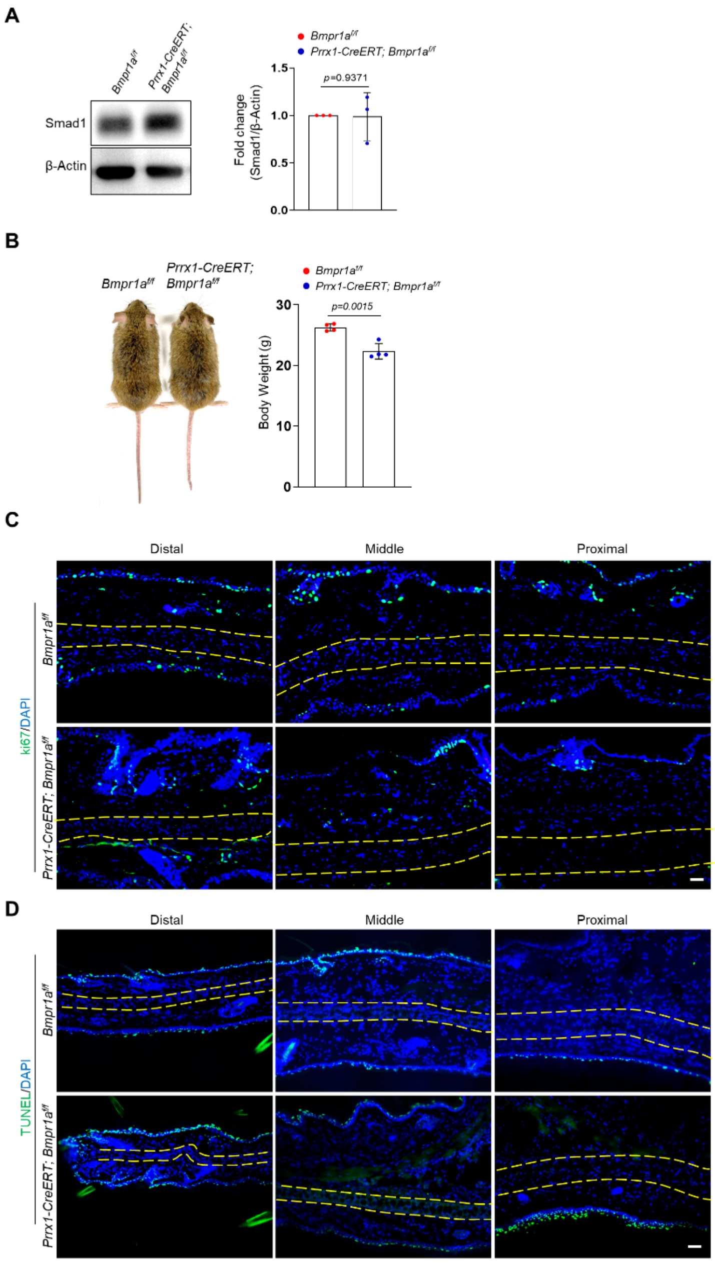


Figure 2-figure supplement 1

**Figure 2-figure supplement 1. The body weight and cell proliferation and apoptosis in the auricle of *Prrx1-CreERT; Bmpr1a$^{f/f}$* mice.**

A. Representative Western blot results of Smad1 expression in the auricle of *Prrx1-CreERT; Bmpr1a$^{f/f}$* and control mice.

B. The size and body weight of *Prrx1-CreERT; Bmpr1a$^{f/f}$* and control mice. N=4.

C. The Ki67 staining results of the ear sections of the mutant and control mice. Scale bars=50 μm.

D. The TUNEL results of the ear sections of the mutant and control mice. Scale bars=50 μm. Unpaired two-tailed Student's t test were applied to evaluate the correlation data in (A and B). $p<0.05$ was considered as statistically significant.

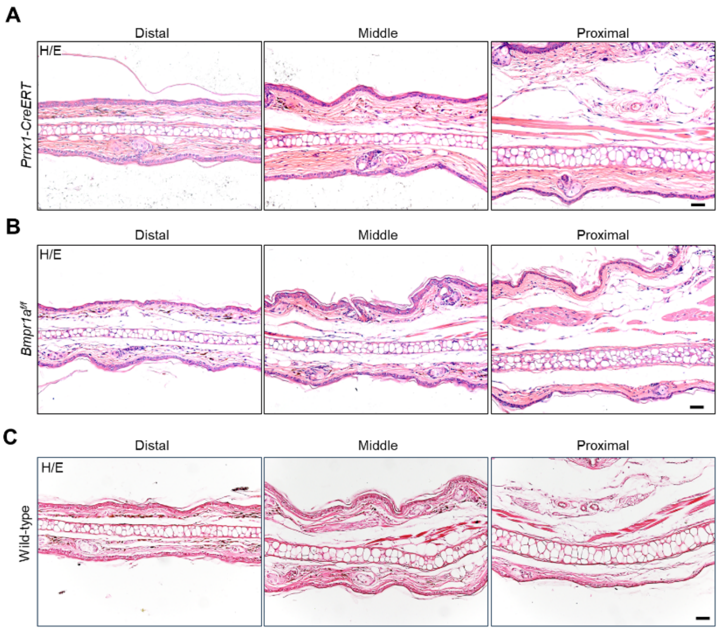


Figure 2-figure supplement 2

**Figure 2-figure supplement 2. No ear phenotype in *Prrx1-CreERT* or *Bmpr1a$^{f/f}$* mice.**

A. No ear phenotype was observed in the *Prrx1-CreERT* (with one *Prrx1* allele deleted) mice. Scale bars=50 µm.

B. No ear phenotype was observed in the *Bmpr1a$^{f/f}$* mice. Scale bars=50 µm.

C. H/E staining results of normal mice. Scale bars=50 µm.

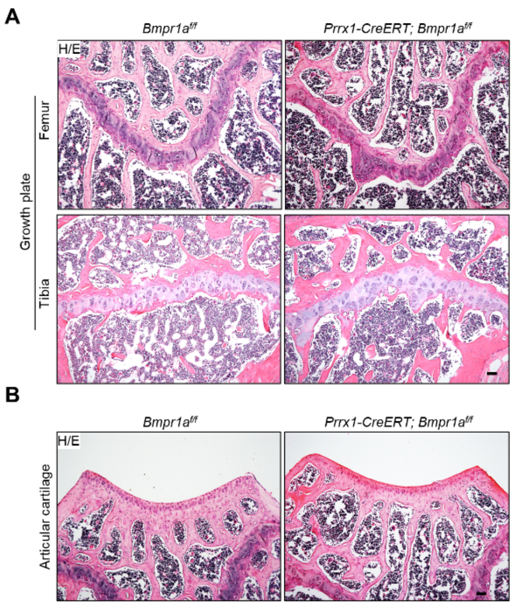


Figure 2-figure supplement 3

**Figure 2-figure supplement 3. Normal growth plate and articular cartilage in adult *Prrx1-CreERT; Bmpr1a*$^{f/f}$ mice receiving TAM.**

A. The epiphysial section of the *Prrx1-CreERT; Bmpr1a*$^{f/f}$ and control mice. Scale bars=100 μm.

B. Articular sections of the *Prrx1-CreERT; Bmpr1a*$^{f/f}$ and control mice. Scale bars=100 μm.

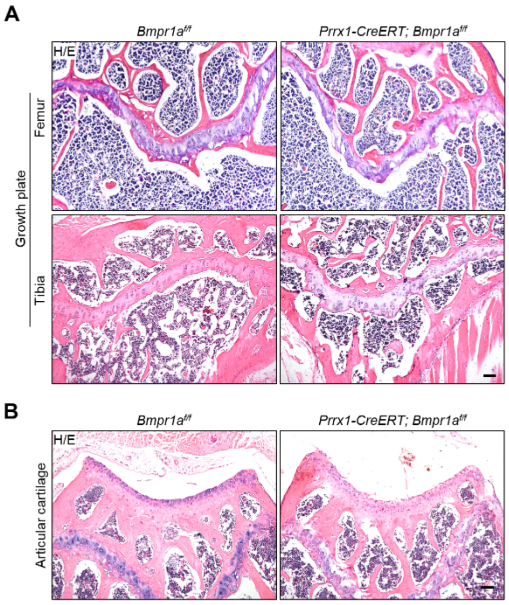


Figure 3-figure supplement 1

**Figure 3-figure supplement 1. Normal growth plate and articular cartilage in *Prrx1-CreERT; Bmpr1a$^{f/f}$* mice receiving TAM at P21.**

A. Epiphysial sections of the *Prrx1-CreERT; Bmpr1a$^{f/f}$* and control mice. Scale bars=100 μm.

B. Articular sections of the *Prrx1-CreERT; Bmpr1a$^{f/f}$* and control mice. Scale bars=100 μm.

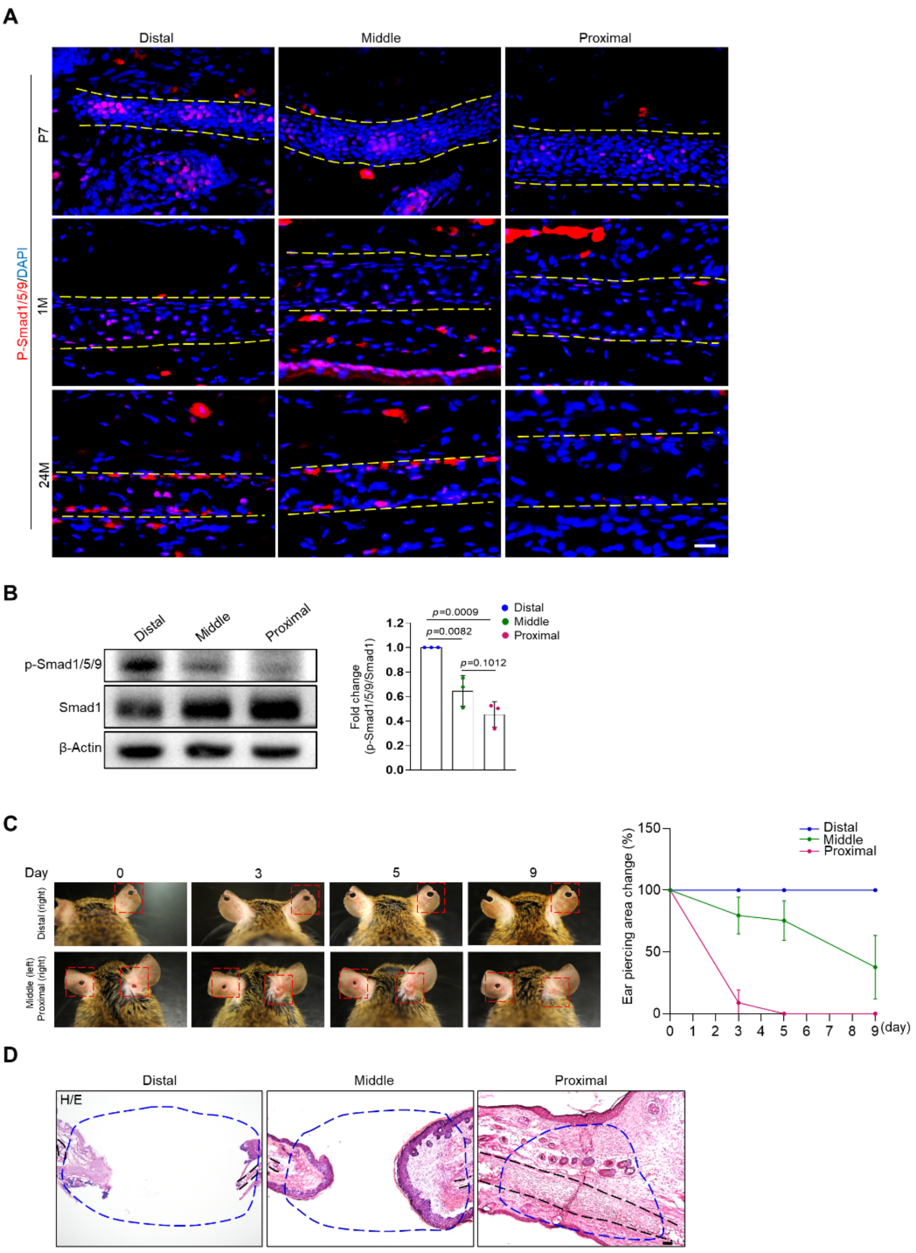


Figure 2-figure supplement 4

**Figure 2-figure supplement 4. Differences in the regenerative activity of the proximal, middle, and distal auricle.**

A. Representative immunostaining results for p-Smad1/5/9 on the proximal, middle, and distal segments of young or adult mice.

B. Western blot analysis of p-Smad1/5/9 in different segments of the auricle. Right panel: quantitation data. N=3.

C. Regeneration of cartilage in different segments of the auricle. Right panel: quantitation data for the diameter of the wounds. N=4.

D. H/E staining results of the regenerative area. Scale bars=100 μm. One-way ANOVA (and nonparametric or mixed) multiple comparisons were applied to evaluate the correlation data in (B and C). $p<0.05$ was considered as statistically significant.

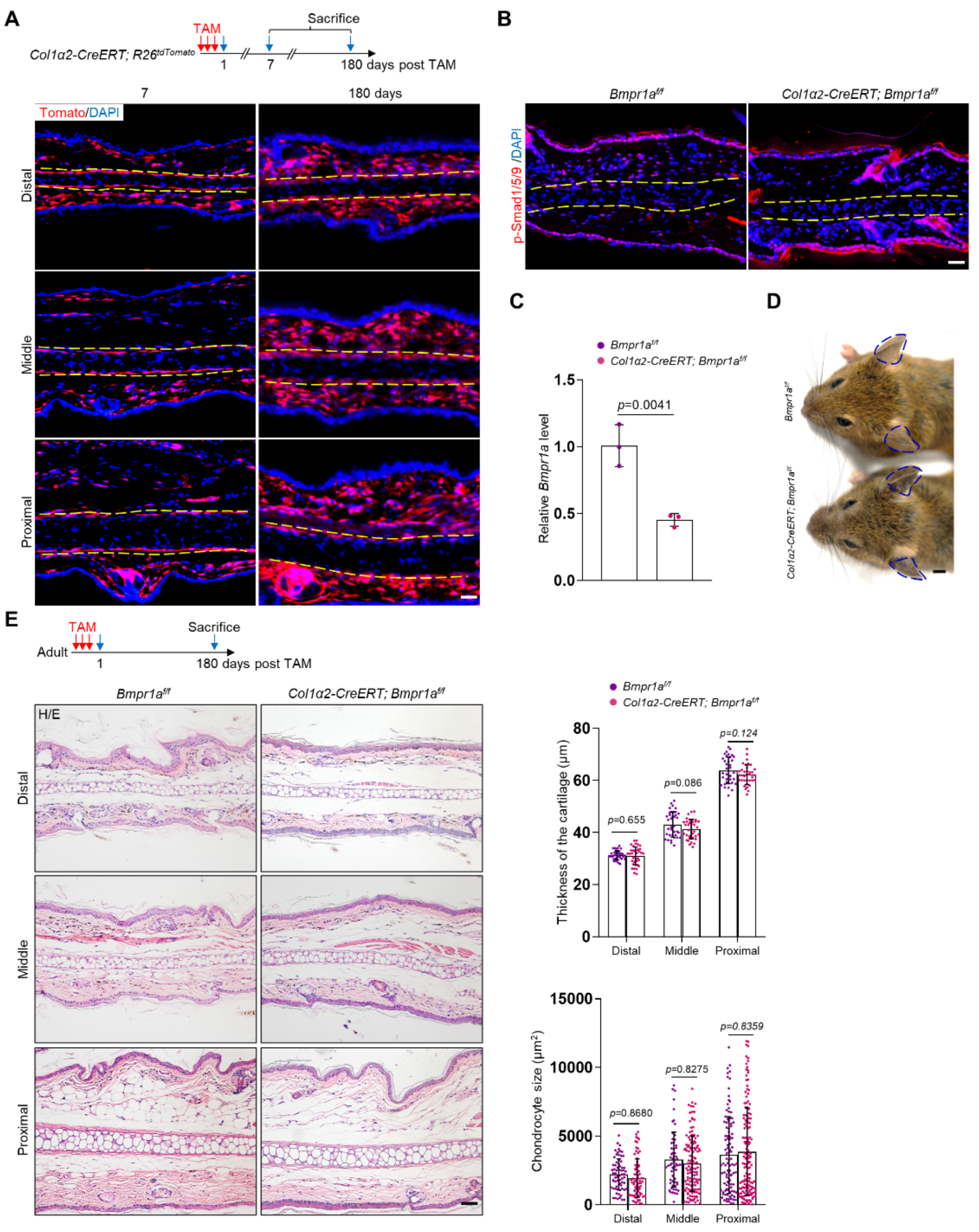


Figure 2-figure supplement 5

**Figure 2-figure supplement 5. Ablation of *Bmpr1a* in dermal cells does not cause microtia.**

A. Genetic tracing experiments showed that *Col1a2* marks all dermal cells on the ear in adult mice. Scale bars=50 μm.

B. Immunostaining of p-Smad1/5/9 on the ear sections of the mutant and control mice. Scale bars=50 μm.

C. qPCR results for *Bmpr1a*. N=3.

D. No ear phenotypes in the *Col1a2-CreERT; Bmpr1a$^{f/f}$* mice.

E. H/E staining of the ear sections of the *Col1a2-CreERT; Bmpr1a$^{f/f}$* mice 180 days after TAM injection. Upper panel: a diagram showing TAM administration and mouse euthanization. Right panels: The thickness of the cartilage and the size of the chondrocytes in the ear of mutant and control mice. Scale bars=50 μm. N=3. Unpaired two-tailed Student's t test were applied to evaluate the correlation data in (C). Two-way ANOVA (or mixed model) multiple comparisons were applied to evaluate the correlation data in (E), $p<0.05$ was considered as statistically significant.

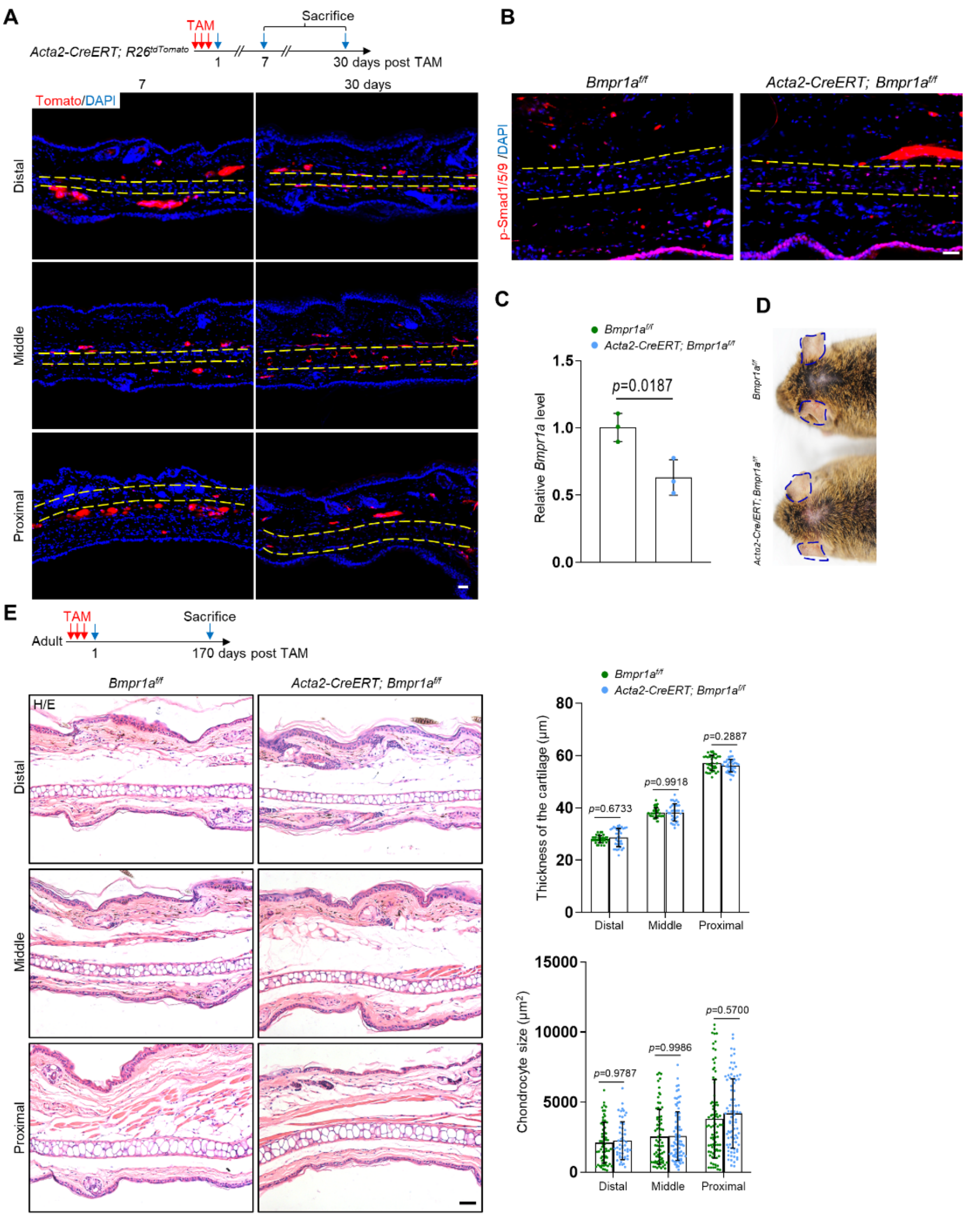


Figure 2-figure supplement 6

**Figure 2-figure supplement 6. Ablation of *Bmpr1a* in smooth muscle cells does not cause microtia**

A. Genetic tracing experiments showed that *Acta2* marks smooth muscle cells and a small portion of chondrocytes in the middle segment of the ear in adult mice. upper panel: a diagram showing TAM administration and mouse euthanization. Scale bars=50 µm.

B. Immunostaining of p-Smad1/5/9 in ear sections of the mutant and control mice. Scale bars=100 µm.

C. qPCR results for *Bmpr1a*. N=3.

D. No ear phenotypes in the *Acta2-CreERT; Bmpr1a$^{f/f}$* mice.

E. H/E staining of the ear sections of *Acta2-CreERT; Bmpr1af/f* mice 170 days after TAM injection. Upper panel: a diagram showing TAM administration and mouse euthanization. Right panels: The thickness of the cartilage and the size of the chondrocytes in the ear of mutant and control mice. Scale bars=50 μm. N=3. Unpaired two-tailed Student's t test were applied to evaluate the correlation data in (C). Two-way ANOVA (or mixed model) multiple comparisons were applied to evaluate the correlation data in (E), $p<0.05$ was considered as statistically significant.

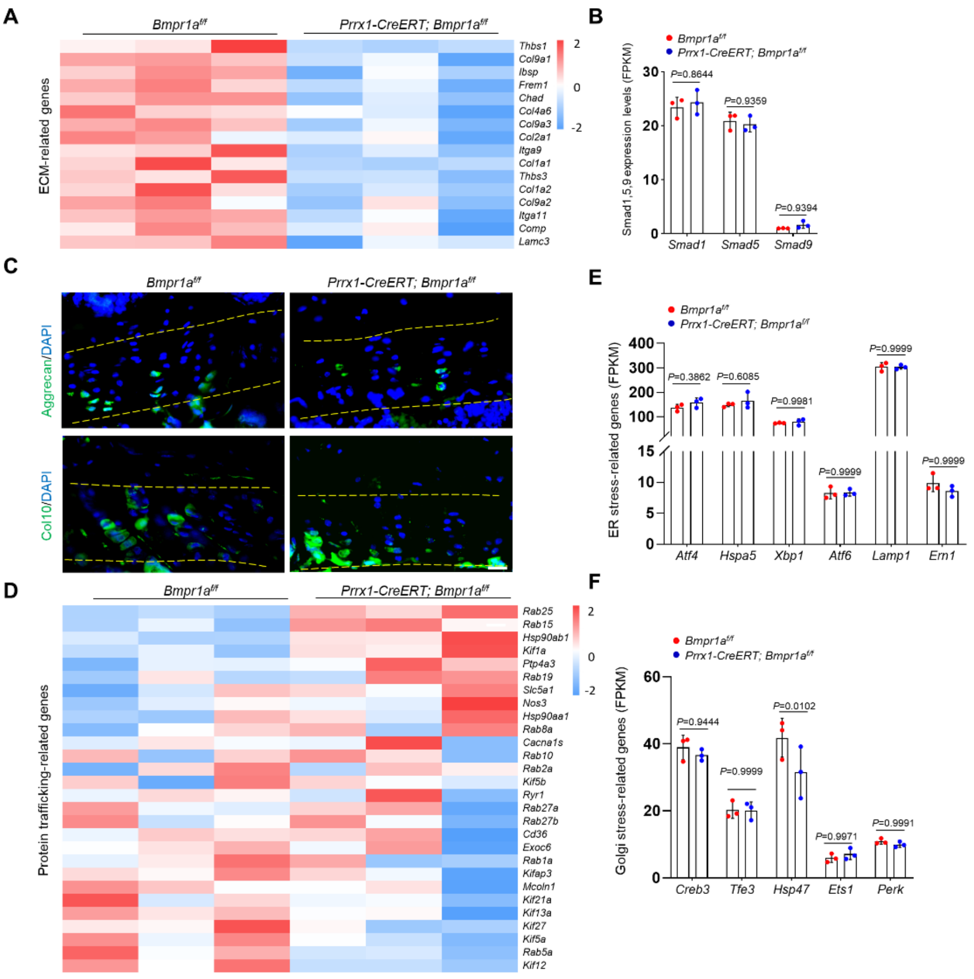


Figure 4-figure supplement 1

**Figure 4-figure supplement 1. Expression of ECM and protein trafficking-related genes in the auricle of control and *Bmpr1a*-deficient mice.**

A. Heatmap of ECM-related genes.

B. FKPM values of Smad1, 5, 9 genes.

C. Representative immunostaining results for Aggrecan and Col10 in growth plates of the mutant and control mice. Scale bars=20 μm.

D. Heatmap of protein trafficking-related genes.

E. FKPM values of ER stress-related genes.

F. FKPM values of Golgi stress-related genes.

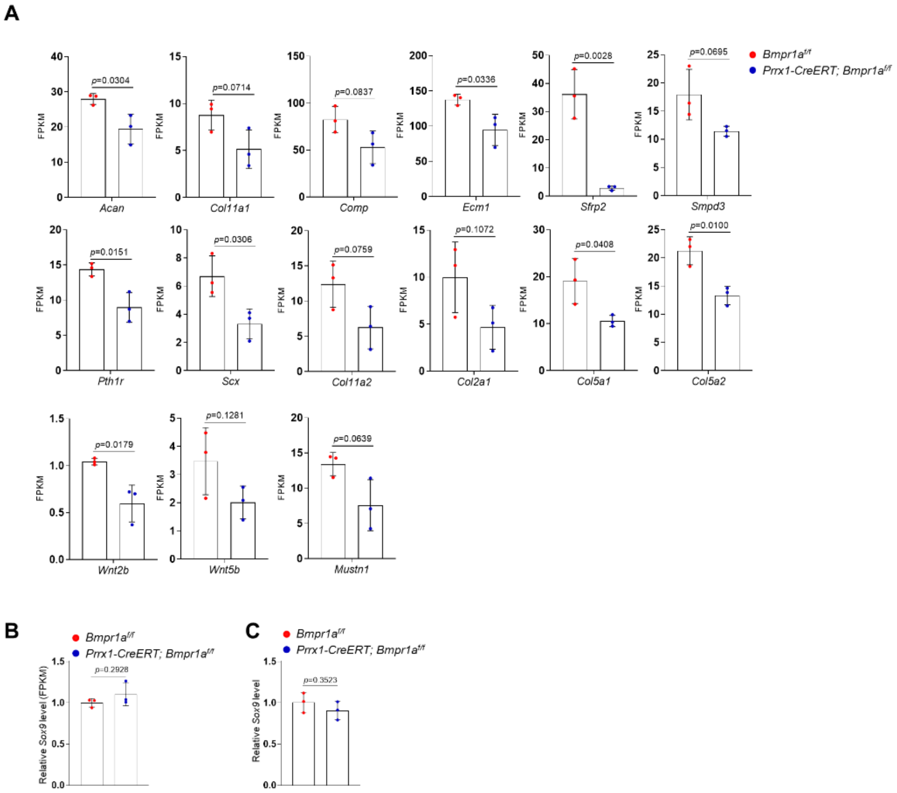


Figure 5-figure supplement 1

**Figure 5-figure supplement 1. Expression of chondrocyte-related genes and *Sox9* in the auricle of control and *Bmpr1a*-deficient mice.**

A. FKPM values for chondrocyte-related genes.

B. FKPM values for Sox9.

C. qPCR results for Sox9.

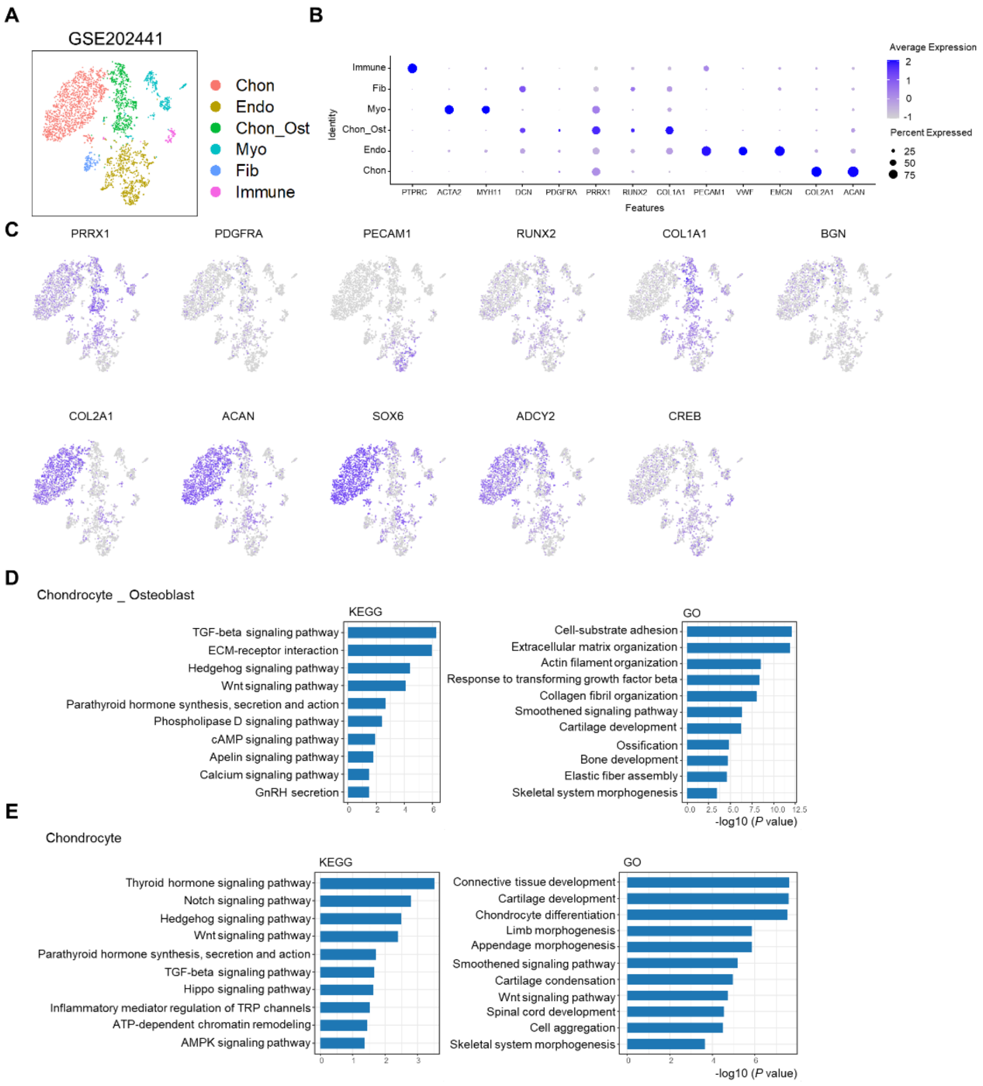


Figure 7-figure supplement 1

**Figure 7-figure supplement 1. Analysis of scRNA-seq data of human microtia pinna samples.**

A. tSNE results of the cell populations in the pinna samples of microtia patients.

B. The expression of marker genes in various cell populations.

C. Analysis of osteoblast and chondrocyte signature genes.

D. KEGG and GO analysis of the osteoblast (Chon_Ost) population.

E. KEGG and GO analysis of the chondrocyte population.
